# Repurposing approach identifies phenylpentanol derivatives as potent azole chemosensitizing agents effective against azole-resistant *Candida* species

**DOI:** 10.1101/696773

**Authors:** Hassan E. Eldesouky, Ehab A. Salama, Xiaoyan Li, Tony R. Hazbun, Abdelrahman S. Mayhoub, Mohamed N. Seleem

## Abstract

The limited number of systemic antifungals and the emergence of azole-resistant *Candida* species constitute a growing challenge to human medicine. Combinatorial drug therapy represents an appealing approach to enhance the activity of, or restore the susceptibility to current antifungals. Here, we evaluated the fluconazole chemosensitization activity of the Pharmakon 1600 drug library against azole-resistant *Candida albicans*. We identified 33 non-antifungal drugs that were able to restore susceptibility to fluconazole in an azole-resistant *C. albicans*. Structural investigation of identified hits revealed phenylpentanol scaffold as a valuable pharmacophore for re-sensitizing azole-resistant *Candida* species to the effect of current azole antifungal drugs. All phenylpentanol derivatives displayed potent fluconazole chemosensitizing activities (ΣFICI 0.13-0.28) and were able to reduce fluconazole’s MIC by 15-31 fold against the tested strain. Particularly pitavastatin displayed the most potent fluconazole chemosensitizing activity (ΣFICI 0.06-0.50). The pitavastatin-fluconazole combination displayed a broad-spectrum synergistic relationship against 90% of the tested strains, including strains of *C. albicans, C. glabrata*, and *C. auris*. Moreover, pitavastatin restored the susceptibility of the multidrug-resistant *C. auris* to the antifungal activities of itraconazole and voriconazole. Additionally, the pitavastatin-fluconazole combination significantly reduced the biofilm-forming abilities of the tested *Candida* species and successfully reduced the fungal burdens in a *Caenorhabditis elegans* infection model. Both pitavastatin and the plain phenylpentanol scaffold were able to interfere significantly with *Candida*’s efflux activities as demonstrated by Nile Red efflux assays and flow cytometry. This study presents phenylpentanol derivatives as potent azole chemosensitizers that warrant further investigation.

## Introduction

Candidiasis is a leading cause of healthcare-associated bloodstream infections and is associated with high morbidities and mortalities. Infections caused by *Candida* species can range from uncomplicated self-limited superficial infections of mucosal membranes to deadly disseminated bloodstream and deep-seated tissue infections often associated with a mortality rate of 42 - 65% [1]. Current epidemiological data portrays *C. albicans* and *C. glabrata* as the major causes of *Candida*-related infections [2-5]. However, the recent emergence of new *Candida* species, such as multidrug-resistant *C. auris* is expected to impact the current epidemiological trends [6-8].

Additionally, *Candida* species are known for their remarkable capabilities of forming robust adherent structures (i.e., biofilms) on surfaces of different abiotic surfaces, such as catheters, and medical implants [9-11]. Biofilms limit the penetration of antifungal drugs and can contribute to treatment failure and chronic infections [12]. Fungal cells residing in biofilms have been reported to have increased expression of efflux genes [13, 14]. Biofilms were also reported to trigger the formation of *Candida* persisters, which can tolerate very high doses of antifungal agents [15]. Collectively, these factors contribute significantly to the remarkable ability of *Candida’s* biofilms to resist the effect of antifungal drugs, especially azoles [16, 17].

Unfortunately, treatment of systemic *Candida* infections is currently limited to only three major drug classes; azoles, polyenes, and echinocandins [18, 19]. Azoles act as inhibitors to the 14α-demethylase enzyme, Erg11, which is vital for ergosterol biosynthesis. Interference with the ergosterol biosynthesis pathway significantly compromises the functions of fungal cell membranes. The limited toxicity, oral bioavailability, and broad-spectrum of antifungal activities made azoles the most commonly prescribed drugs for controlling and treating *Candida* infections [19, 20]. The high dependence of clinicians on azole antifungal agents has been associated with the emergence of azole-resistant *Candida* strains [21, 22].

Given the clinical importance of azole antifungals, there is a pressing need for potent co-drugs that would augment the antifungal effect of azole drugs, particularly against *Candida* biofilms and azole-resistant strains. In this study, we explored the azole chemosensitizing activity of ∼1600 approved drugs and clinical molecules from the Pharmakon drug library. We identified 33 non-antifungal drugs that were able to restore susceptibility to fluconazole in an azole-resistant *C. albicans* strain. A more in-depth structural investigation of identified hits revealed the phenylpentanol scaffold as a valuable pharmacophore for re-sensitizing azole-resistant *Candida* species to the effect of current azole antifungal drugs. The most potent phenylpentanol derivative (pitavastatin) was further investigated in combination with azole drugs against strains of *C. albicans, C. glabrata*, and the multidrug-resistant *C. auris* was evaluated for the ability to inhibit *Candida* biofilm formation and was assessed for the ability to reduce *Candida* burdens in infected nematodes. Furthermore, the effect of pitavastatin and the plain phenylpentanol scaffold on the efflux activities of *Candida* strains with known efflux mechanisms was evaluated.

## Results

### Identification of drugs with fluconazole chemosensitizing activity

The Pharmakon 1600 drug library was initially screened at 16 µM concentration against *C. albicans* NR-29448, a strain that displayed high-level resistance to several azole antifungal drugs including fluconazole, itraconazole, and voriconazole. In order to explore the azole chemosensitizing activities of the Pharmakon 1600 drug library, two rounds of screening were performed either in the presence or absence of a sub-inhibitory concentration of fluconazole (32 µg/ml). This high concentration of fluconazole was selected to maximize the initial pool of active hits. Drugs that inhibited the growth of *C. albicans*, only in the presence of fluconazole, were identified as “positive hits.”. Our initial screening identified 33 drugs that exhibited synergistic interactions with fluconazole against the azole-resistant strain *C. albicans* NR-29448 (Table-1). The initial screening identified novel fluconazole chemosensitizers that have never been reported before, such as bufexamac, apomorphine, and sulfaquinoxaline. As expected, several drugs with reported fluconazole chemosensitization activity were identified, such as the calcineurin inhibitors cyclosporin, tacrolimus, sirolimus, nisoldipine, benazepril, and estrogen receptor modulators [23-25]. Also, several antibacterial agents with reported fluconazole chemosensitization activities were identified, such as sulfa drugs, doxycycline, and clofazimine [25-27]. However, structural analysis of the identified hits revealed a common phenylpentanol core present in many of these structurally diverse hit compounds. Phenylpentanol, or its saturated cyclohexylpentanol, was observed in nine drugs, namely; apomorphine, atorvastatin, lovastatin, simvastatin, cholecalciferol, doxycycline, quinestrol, testosterone, and norgestimate (Figure-1).

**Table 1.**
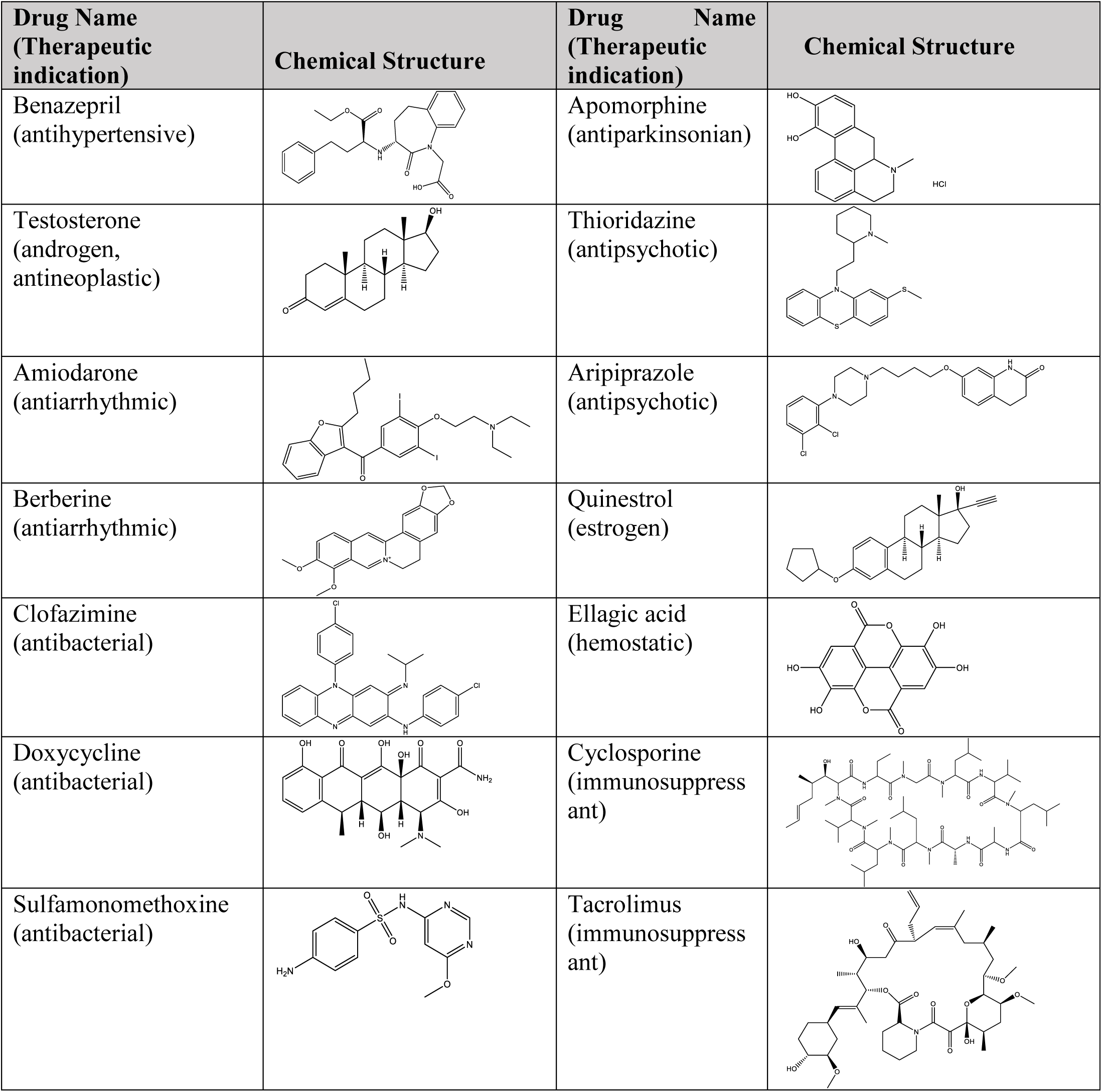
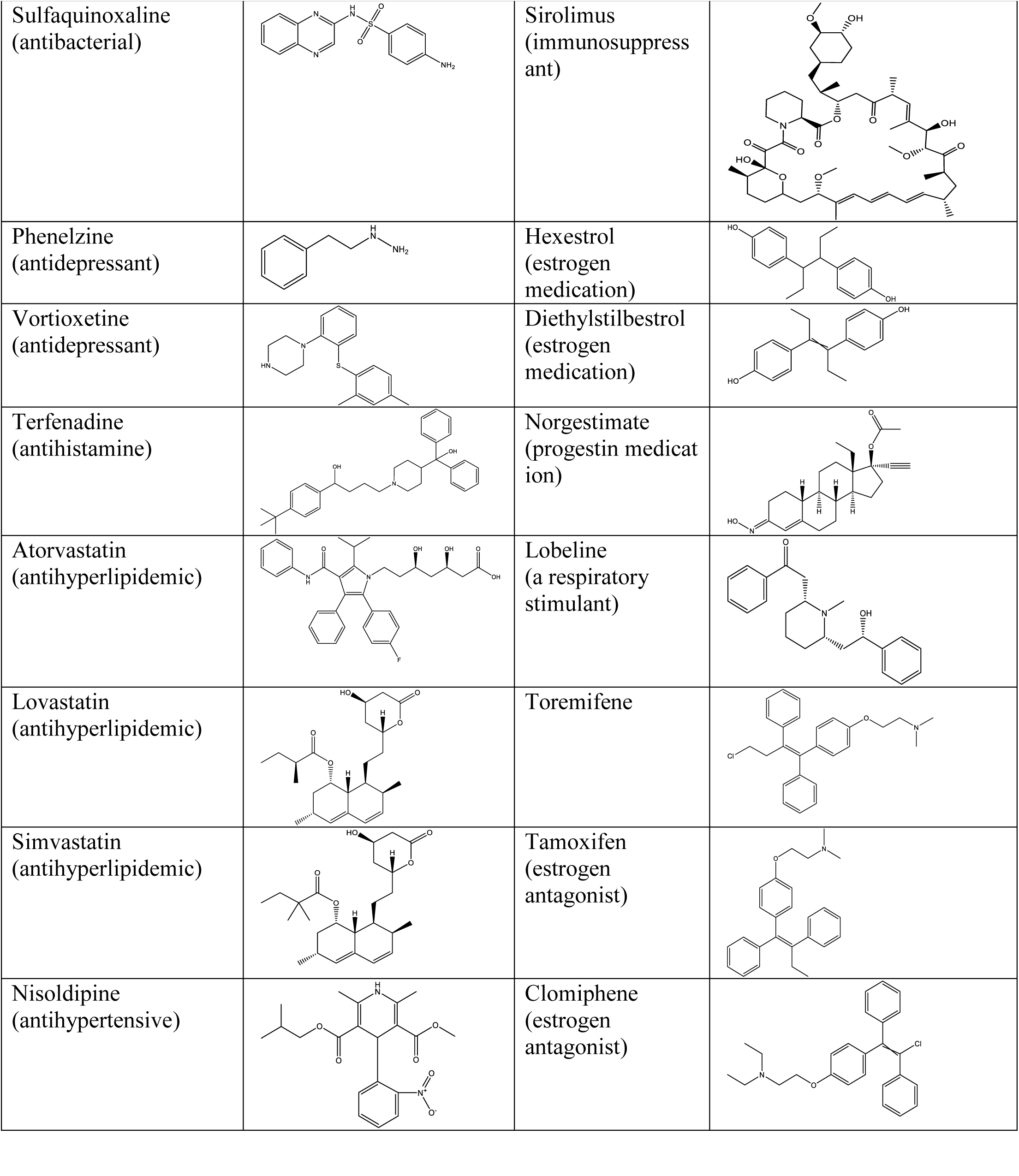
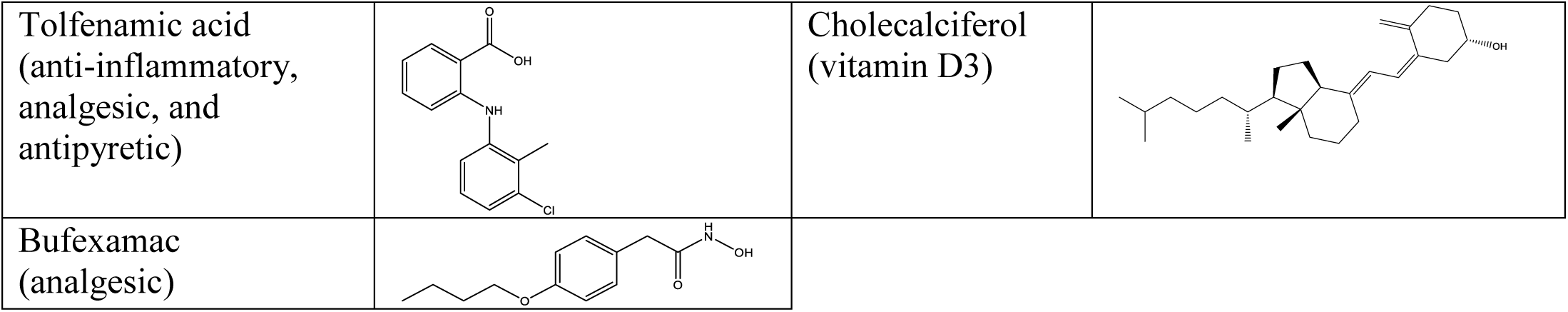
Names, therapeutic indications, and chemical structures of positive hit compounds identified from screening the fluconazole chemosensitizing activities of the Pharmakon drug library against *C. albicans* NR-29448.

**Figure 1.**
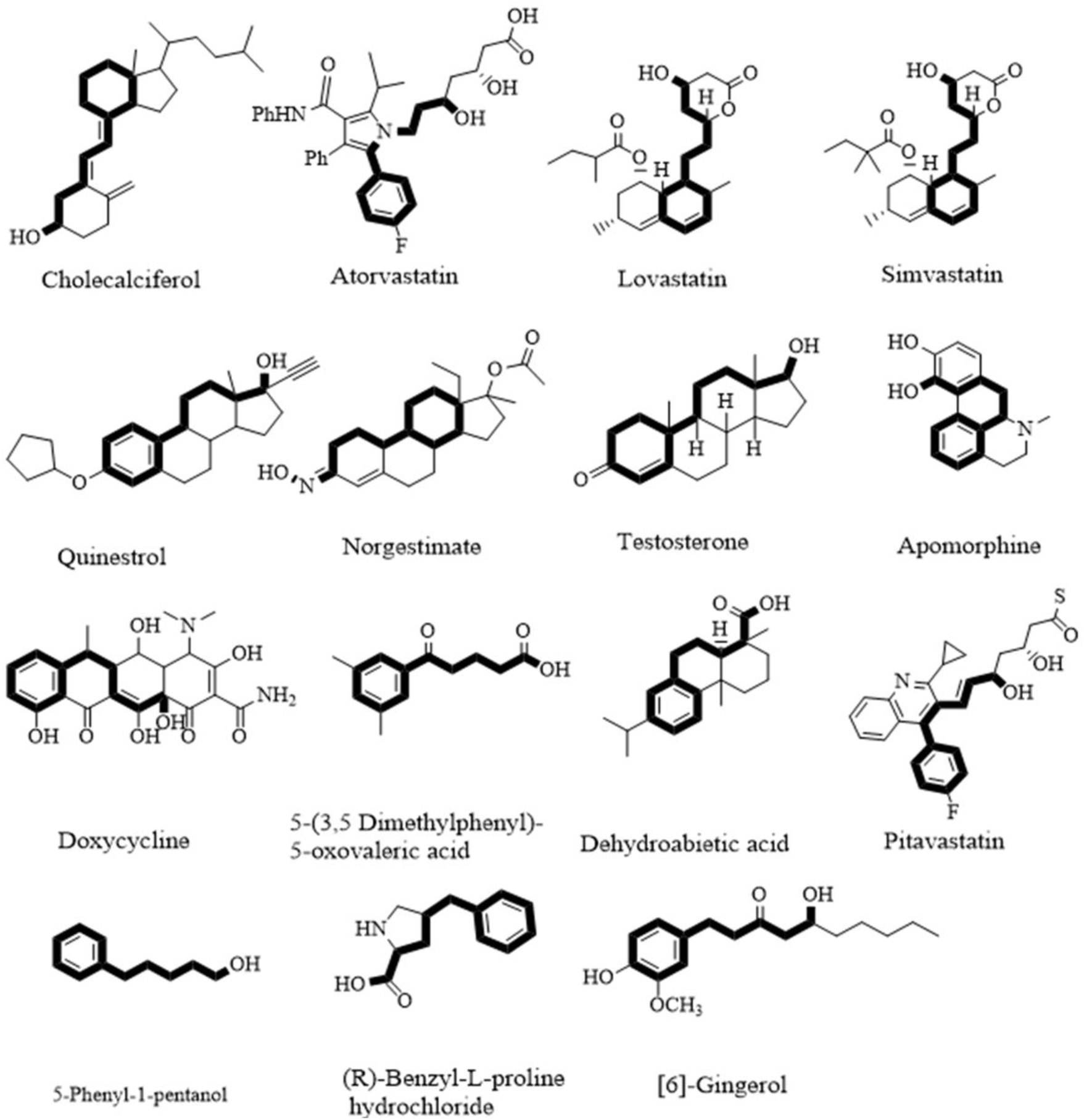
Chemical structures of drugs that contain phenylpentanol and its closely related pharmacophores (highlighted in bold black bonds); stereochemistry was not considered for the sake of clarity

### Synergistic interactions between fluconazole and phenylpentanol derivatives against *C. albicans* NR-29448

Microdilution checkerboard assays were used to assess the synergistic interactions between phenylpentanol derivatives and fluconazole against the azole-resistant *C. albicans* NR29448 strain. Expectedly, as shown in Table-2, all phenylpentanol derivatives displayed potent fluconazole chemosensitizing activities (ΣFICI = 0.13-0.28) and were able to reduce fluconazole’s MIC by 15-31 fold against the tested strain. To confirm that the phenylpentanol core is vital for the observed azole chemosensitizing activities, we evaluated the fluconazole chemosensitizing activities of six additional phenylpentanol derivatives (entries 10-15 in Table-2) that were not present in the Pharmakon drug library. Notably, four additional phenylpentanol compounds, namely, 5-phenylpentanol, oxovaleric acid, dehydroabietic acid, and pitavastatin, exhibited strong synergistic interactions with fluconazole against *C. albicans* NR29448. Interestingly, pitavastatin displayed the most potent fluconazole chemosensitizing activity (ΣFICI = 0.06) and was superior to all the tested phenylpentanol derivatives, including the pharmacologically-related statins, which were reported to have fluconazole-chemosensitizing activities [28, 29]. Pitavastatin, at 0.25 µg/ml, was able to reduce the MIC of fluconazole by 31-fold against *C. albicans* NR29448. Due to its potent fluconazole chemosensitizing activity, pitavastatin was selected for subsequent experimental investigation.

**Table 2.**
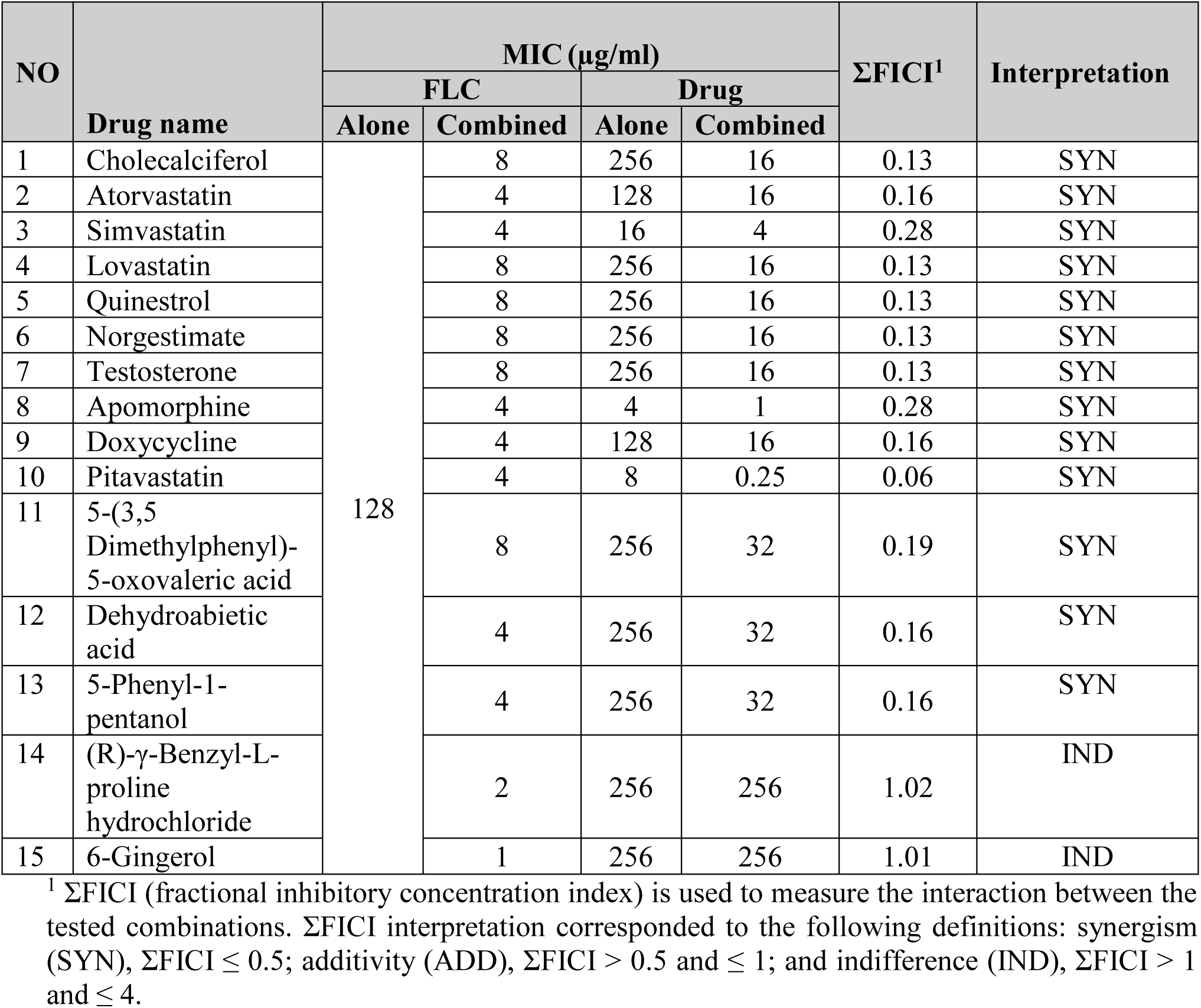
Effect of different combinations of phenylpentanol derivatives and fluconazole (FLC) against *C. albicans* NR-29448.

### Pitavastatin displays potent azole chemosensitizing activity against different *Candida* species

After identifying pitavastatin as the most potent azole-chemosensitizing agent from the compounds/drugs screened against *C. albicans* NR-29448, we evaluated if its synergistic interactions with fluconazole would extend to other strains and species of *Candida*. As shown in Table-3, pitavastatin exhibited a synergistic relationship with fluconazole (ΣFICI = 0.06-0.50) against 16, out of 18 *Candida* strains (89%), resulting in significant reductions in fluconazole’s MIC values (by 3-31 folds). Only *Candida albicans* strains TWO7241 and SC-MRR1^R34A^ did not respond to the synergistic relationship between pitavastatin and fluconazole.

**Table 3.** Effect of pitavastatin-fluconazole (FLC) combination against different *Candida* strains.

| Fungal Strain | MIC (µg/ml) |  |  |  | ΣFICI<sup>1</sup> | Interaction |
| --- | --- | --- | --- | --- | --- | --- |
|  | FLC |  | Pitavastatin |  |  |  |
|  | Alone | Combined | Alone | Combined |  |  |
| *C. albicans* SC5314 | 0.125 | 0.0312 | 4 | 1 | 0.50 | SYN |
| *C. albicans* NR-29448 | > 64 | 4 | 8 | 0.25 | 0.06 | SYN |
| *C. albicans* NR-29437 | > 64 | 4 | 8 | 0.5 | 0.09 | SYN |
| *C. albicans* ATCC 26790 | > 64 | 4 | 8 | 0.5 | 0.09 | SYN |
| *C. albicans* ATCC MYA-573 | > 64 | 4 | 16 | 4 | 0.28 | SYN |
| *C. albicans* TWO7241 | 32 | 16 | 8 | 4 | 1.00 | IND |
| *C. albicans* TWO7243 | 64 | 16 | 64 | 4 | 0.31 | SYN |
| *C. albicans* SC-TAC1<sup>R34A</sup> | 2 | 0.25 | 32 | 4 | 0.25 | SYN |
| *C. albicans* SC-MRR1<sup>R34A</sup> | 2 | 1 | 8 | 2 | 0.75 | ADD |
| *C. glabrata* ATCC 66032 | 4 | 1 | 64 | 16 | 0.50 | SYN |
| *C. glabrata* ATCC MYA-2950 | 8 | 1 | 64 | 16 | 0.38 | SYN |
| *C. glabrata* ATCC 2001 | 4 | 0.5 | 64 | 16 | 0.38 | SYN |
| *C. glabrata* HM-1123 | 4 | 1 | 64 | 16 | 0.50 | SYN |
| *C. auris* 385 | 256 | 32 | 128 | 32 | 0.38 | SYN |
| *C. auris* 386 | 256 | 64 | 128 | 16 | 0.38 | SYN |
| *C. auris* 388 | 256 | 64 | 64 | 16 | 0.50 | SYN |
| *C. auris* 389 | 256 | 32 | 64 | 8 | 0.25 | SYN |
| *C. auris* 390 | 256 | 16 | 64 | 16 | 0.31 | SYN |

<sup>1</sup> ΣFICI (fractional inhibitory concentration index) is used to measure the interaction between the tested combinations. ΣFICI interpretation corresponded to the following definitions: synergism (SYN), ΣFICI ≤ 0.5; additivity (ADD), ΣFICI > 0.5 and ≤ 1; and indifference (IND), ΣFICI > 1 and ≤ 4.

Next, pitavastatin was evaluated in combination with different azoles, including voriconazole and itraconazole, against the tested *Candida* strains. Similar to its effect with fluconazole, pitavastatin possessed a broad-spectrum synergistic relationship with voriconazole against 16 strains of *C. albicans, C. glabrata*, and *C. auris* (ΣFICI ranged from 0.15–0.50, Supplementary Table-2). However, the combination of pitavastatin and itraconazole displayed a narrow-spectrum synergistic relationship, as only 9 out of 18 of the tested *Candida* strains (50%) responded to the pitavastatin-itraconazole combination (Supplementary Table-3).

### Effect of the pitavastatin-fluconazole combination on the growth kinetics of *Candida* species

The checkerboard microdilution assays revealed a significant synergistic relationship between pitavastatin and fluconazole that was effective against *C. albicans, C. glabrata*, and *C. auris*. To confirm this relationship, we monitored the effect of the pitavastatin-fluconazole combination on growth kinetics of the three most sensitive strains from each *Candida* species investigated. As shown in Figure-2, combinations of pitavastatin and fluconazole (at concentrations corresponding to the MIC values obtained when combined in the checkerboard assay) significantly inhibited the growth of *C.* albicans NR-29448 (panel A), *C. glabrata* MYA-2950 (panel B), and *C. auris* 390 (panel C). However, neither pitavastatin nor fluconazole alone (at the same concentrations used when tested in combination) was able to inhibit the growth of *Candida* strains significantly.

**Figure 2.**
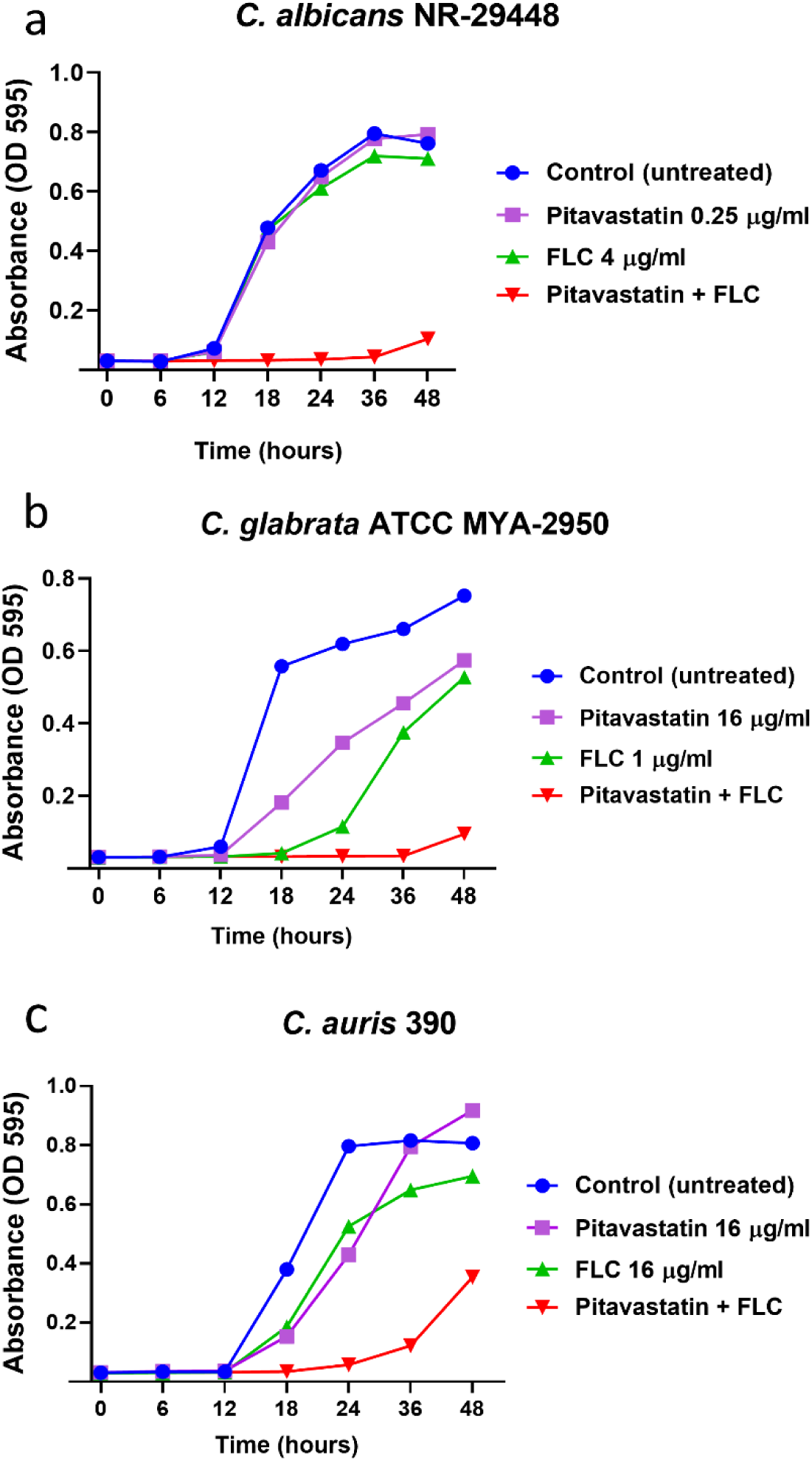
Effect of the pitavastatin-fluconazole combination on the growth kinetics of different *Candida* species. Overnight cultures of a) *C. albicans* NR-29448, b) *C. glabrata* ATCC MYA-2950, and c) *C. auris* 390 were diluted to 0.5-2.5 × 10^3^ CFU/ml in RPMI 1640 medium. Cells were treated with pitavastatin, fluconazole (FLC), or a combination of the two drugs at the indicated concentrations. Cells were incubated at 35 °C for 48 h, and OD_595_ values were measured at different time points (0, 6, 12, 18, 24, 36 and 48 h).

### The pitavastatin-fluconazole combination significantly reduces the biofilm-forming abilities of *Candida* species

We next investigated whether the synergistic relationship between azole drugs and pitavastatin could interfere with the biofilm-forming ability of *Candida*. Compared to single treatments with either fluconazole or pitavastatin, incubating the tested *Candida* species with pitavastatin (at 0.5 × MIC) in the presence of a subinhibitory concentration of fluconazole (2 µg/ml) resulted in a significant reduction in the biofilm-forming abilities of *C. albicans* NR-29448 (73% reduction, Figure-3, panel a), *C. glabrata* HM-1123 (50% reduction, Figure-3, panel b), and *C. auris* 385 (40% reduction, Figure-3, panel c).

**Figure 3.**
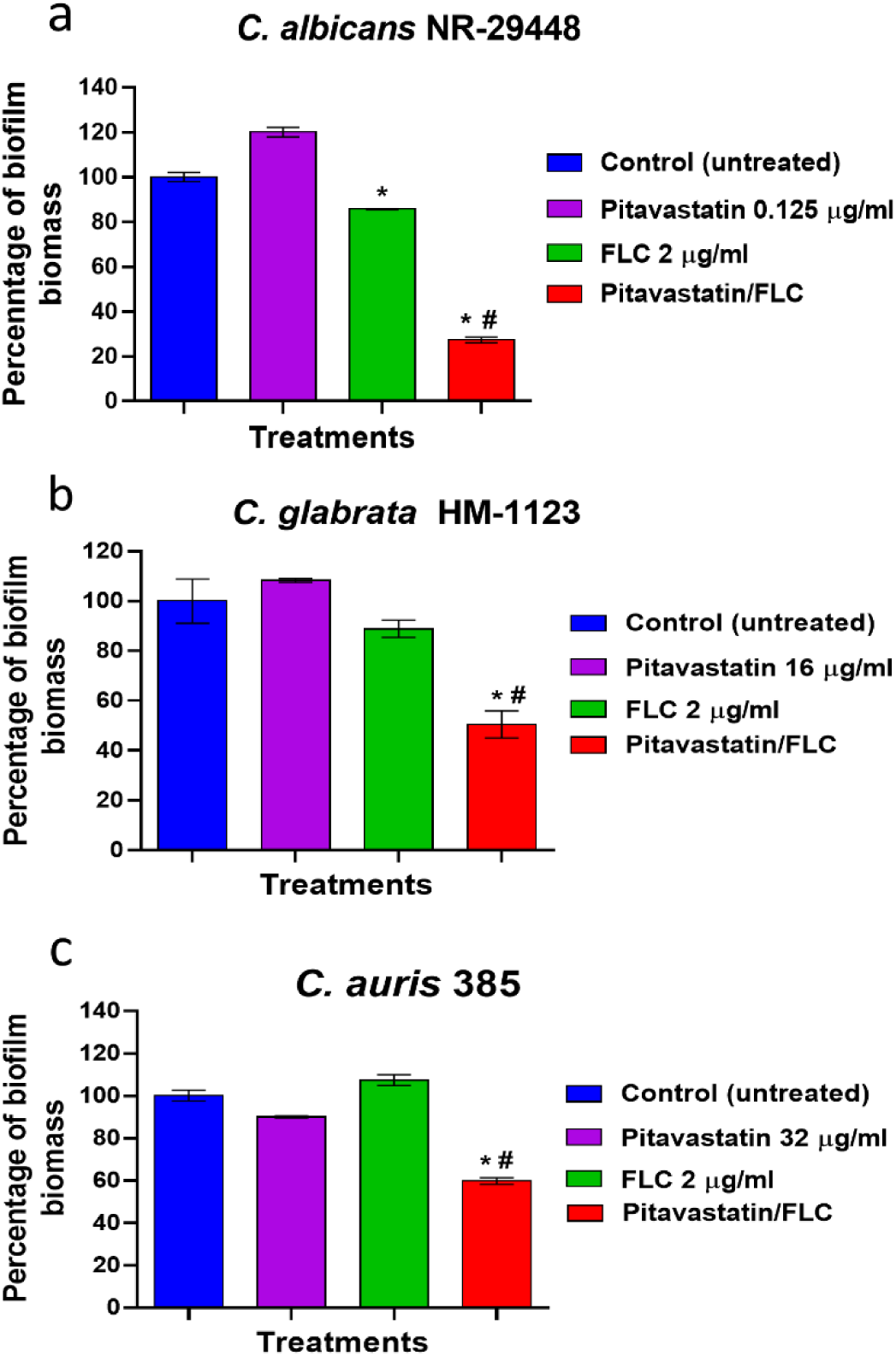
Anti-biofilm activity of the pitavastatin-fluconazole combination. The effect of the pitavastatin-fluconazole (FLC) combination was tested on the biofilm-forming ability of a) *C. albicans* NR-29448, b) *C. glabrata* HM-1123, and c) *C. auris* 385, respectively. Fresh overnight cultures of the tested *Candida* strains were diluted 1:100 in RPMI 1640 medium. Cells were treated with pitavastatin, fluconazole (FLC), or a combination of the two drugs, at the indicated concentrations. Yeast strains were then incubated at 35 ^°C^ for 24 h. The absorbance of crystal violet-stained biofilms was measured at OD_595_. * indicates a significant difference between each treatment compared to the untreated control (*P* < 0.05, as determined by one-way ANOVA with post-hoc Dunnet’s test for multiple comparisons). Whereas # indicates a significant difference between the tested pitavastatin-fluconazole combination relative to the single treatment with either fluconazole or pitavastatin (*P* < 0.05, as determined by one-way ANOVA with post-hoc Dunnet’s test for multiple comparisons).

### Pitavastatin significantly interferes with the ABC-mediated efflux activity of *Candida*

Nile red is a known substrate for the two major types of membrane transporters (ABC and MFS) expressed by *Candida* [30]. Therefore Nile red was used to study the effect of pitavastatin, and 5-phenylpentanol on the efflux activity of *C. albicans* strains with known mechanisms of efflux hyperactivity (SC-TAC1^R34A^, TWO7243, SC-MRR1^R34A^, and TWO7241). Using the Nile Red efflux assay and flow cytometric analysis, we studied the effect of pitavastatin on *Candida’s* efflux activity against two major membrane exporters, the ATP-binding cassette (ABC) and the major facilitator superfamily (MFS) membrane transporters. As shown in Figure-4, pitavastatin (at 0.25 × MIC) significantly reduced Nile Red fluorescence intensity in strains SC-TAC1^R34A^ (panel a) and TWO7243 (panel B). However, the Nile Red fluorescence intensity remained unaffected in strains SC-MRR1^R34A^ (panel c) and TWO7241 (panel d).

**Figure 4:**
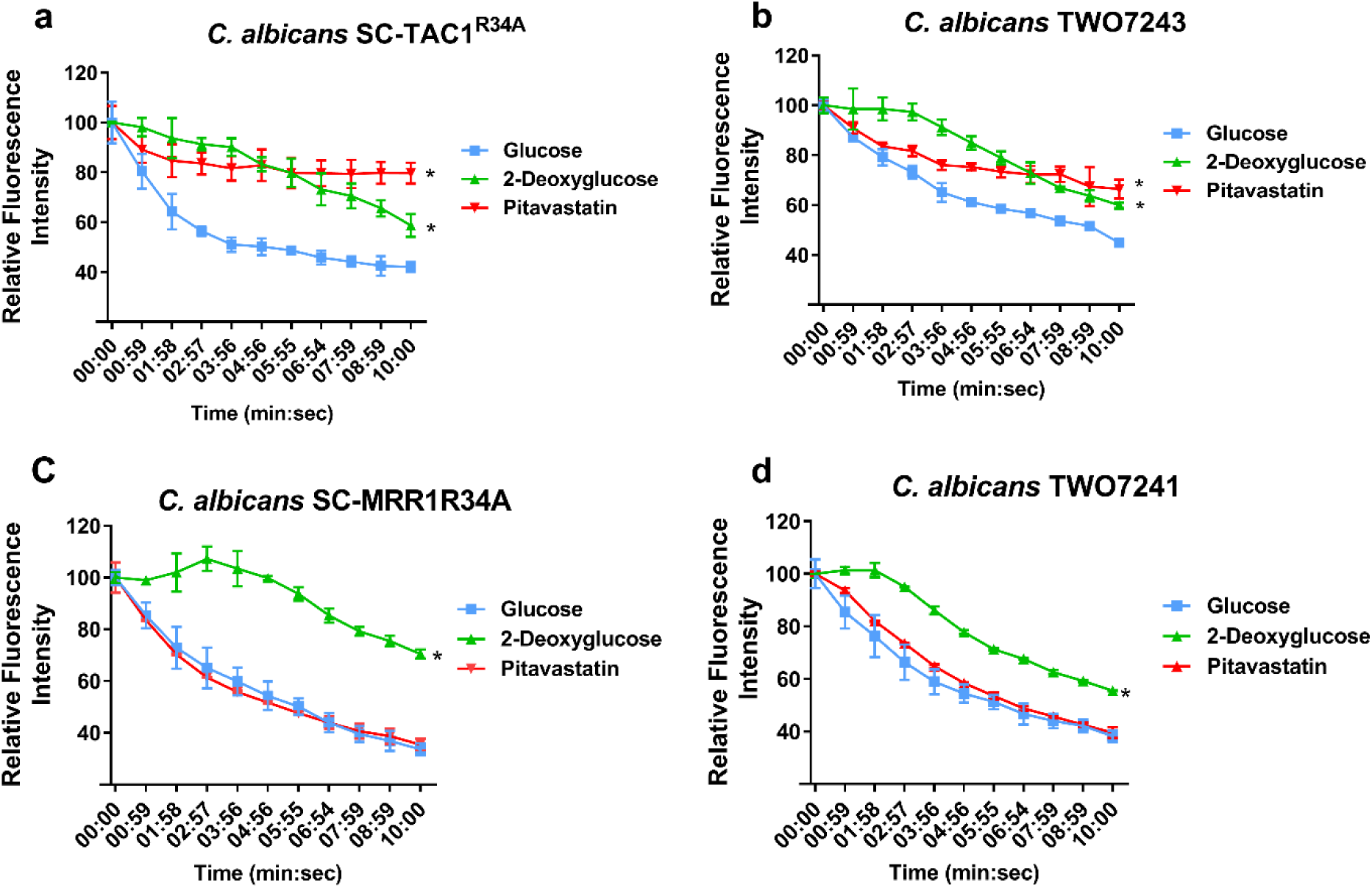
Effect of pitavastatin on Nile red efflux by different efflux hyperactive *Candida* strains. The effect of pitavastatin on Nile Red efflux by the ABC efflux hyperactive strains a) SC-TAC1R34A and b) TWO7243. The effect of pitavastatin on Nile Red efflux mediated by the MFS efflux hyperactive strains c) SC-MRR1R34A and d) TWO7241. Efflux-activated strains were starved overnight in PBS and were loaded with the fluorescent dye, Nile red (7.5 µM). Pitavastatin was added to the yeast cells at a sub-inhibitory concentration (0.25 × MIC). Efflux of Nile red was initiated by adding glucose (final concentration 40 mM) to all treatments at time zero (00:00), except for the 2-deoxyglucose treatment. The Nile red fluorescence intensity was then monitored over 10 minutes and is expressed as the percentage of change in the fluorescence intensity. * indicates a significant difference between treatment with either 2-deoxyglucose or pitavastatin compared to the glucose treated control (*P* < 0.05, as determined by multiple t-tests using Holm-Sidak statistical method for multiple comparisons).

The effect of pitavastatin on *Candida’s* efflux activities was further confirmed using flow cytometry. *C. albicans* SC-TAC1^R34A^ exhibited a significant increase in Nile Red fluorescence intensity when exposed to pitavastatin (at 0.25 × MIC), compared to the glucose treated control (Figure 5a). The pitavastatin effect on Nile Red efflux observed in strain SC-TAC1^R34A^ was equivalent to the effect of 2-deoxyglucose, which interferes with *Candida*’s efflux activities by hindering its ability to produce ATP. However, pitavastatin did not affect the Nile Red efflux activity of *C. albicans* SC-MRR1^R34A^ and the Nile red fluorescence intensity was similar to the glucose treated control (Figure 5b), which is in agreement with our previous observations.

**Figure 5.**
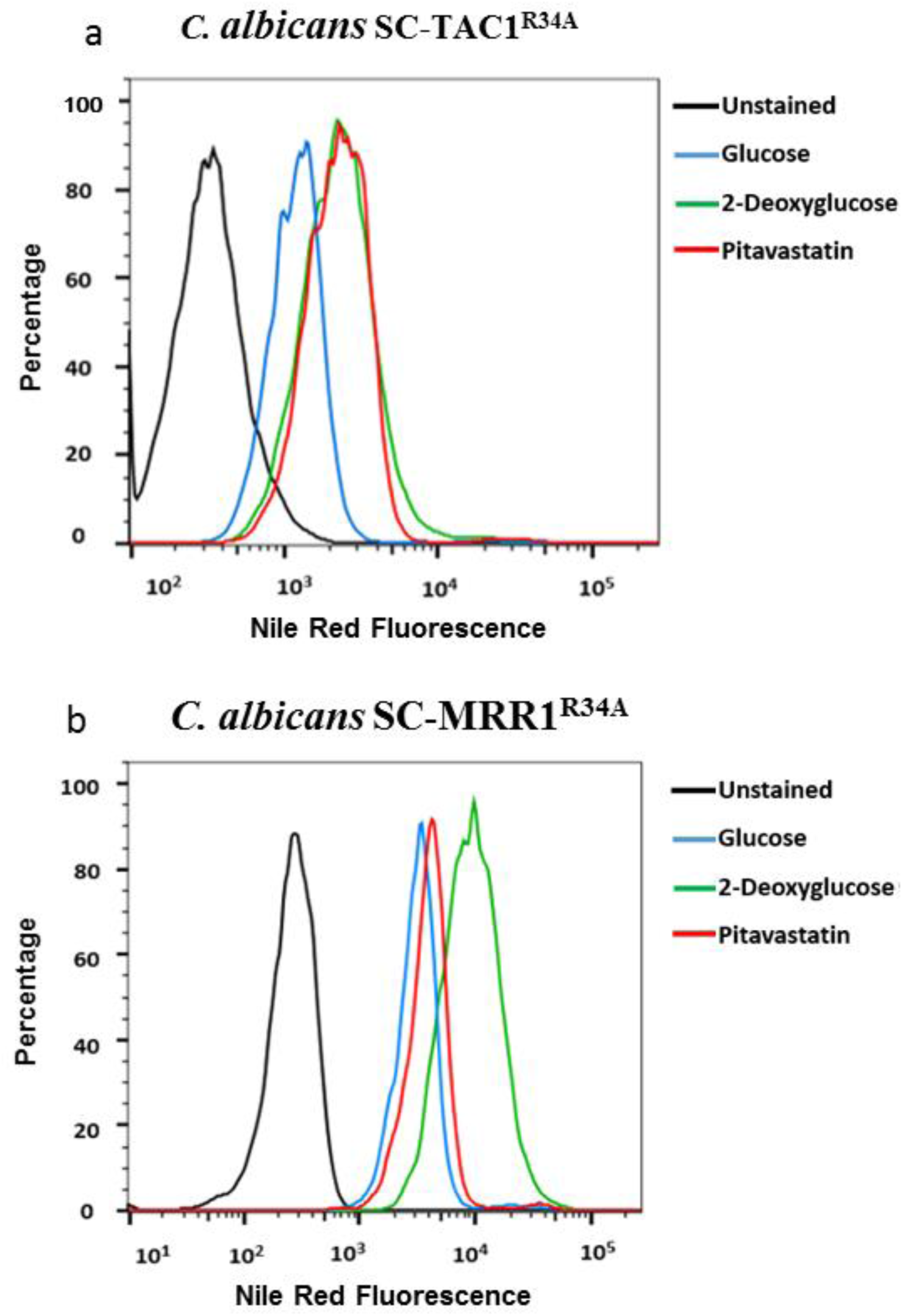
Flow cytometry analysis of Nile red efflux from two *C. albicans* strains treated with pitavastatin. Flow cytometric analysis of the effect of pitavastatin (at 0.25 × MIC) on Nile red efflux from a) *C. albicans* SC-TAC1^R34A^ and b) *C. albicans* SC-MRR1^R34A^. Histograms represent overlaid flow cytometry data as a percentage of unstained or Nile Red-stained cells treated with either glucose, 2-deoxyglucose, or pitavastatin (0.25 µg/ml). The Nile Red fluorescence intensity was recorded at a single time point (30 minutes following pitavastatin treatment).

### Effect of the pitavastatin-fluconazole combination on reducing the fungal burden in *Caenorhabditis elegans* infected with azole-resistant *Candida*

In order to validate our *in vitro* results, *C. elegans* was utilized as an animal model to investigate the fluconazole chemosensitizing activity of pitavastatin. Compared to individual treatments with either fluconazole or pitavastatin, the pitavastatin-fluconazole combination significantly reduced the mean fungal CFU burden of infected nematodes by approximately 97% (against *C. albicans* NR-29448, Figure 6a), 68% (against *C. glabrata* ATCC MYA-2950, Figure 6b), and 34% (against *C. auris* 390, Figure 6c).

**Figure 6.**
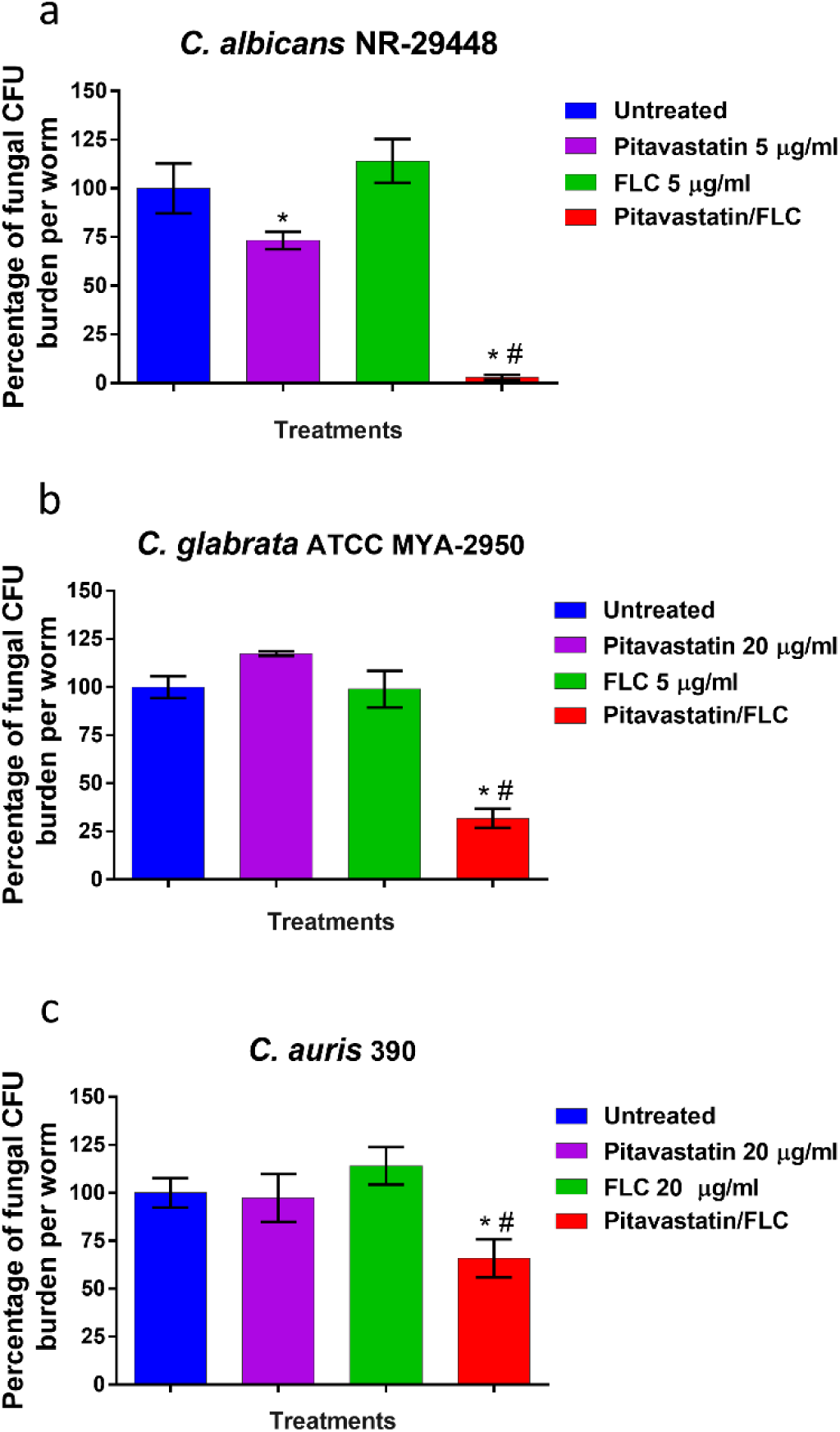
*In vivo* efficacy of the pitavastatin-fluconazole combination in reducing fungal burden in infected *C. elegans*. *C. elegans* strain AU37 genotype [glp-4(bn2) I; sek-1(km4) X], was co-incubated with yeast suspensions of a) *C. albicans* NR-29448, b) *C. glabrata* ATCC MYA-2950, or c) *C. auris* 390 using an inoculum size of ∼5 × 10^7^ CFU/ml for three hours at room temperature. Infected nematodes were washed with PBS and then treated with the pitavastatin-fluconazole combination at the respective concentration. Treatment with PBS, pitavastatin alone, or fluconazole alone served as controls. After 24 h of treatment, worms were lysed to determine the fungal burden (CFU/worm) after treatment. * indicates a significant difference between each treatment compared to the untreated control (*P* < 0.05, as determined by one-way ANOVA with post-hoc Dunnet’s test for multiple comparisons). Whereas # indicates a significant difference between the tested pitavastatin-fluconazole combination relative to the single treatment with either fluconazole or pitavastatin (*P* < 0.05, as determined by one-way ANOVA with post-hoc Dunnet’s test for multiple comparisons).

## Discussion

In order to explore new co-drugs capable of sensitizing resistant strains of *Candida* to the effect of azoles, we investigated the azole chemosensitizing activity of 1600 drugs and clinical molecules present in the Pharmakon drug library. Although the initial screening revealed a set of structurally diverse molecules, we observed the phenylpentanol scaffold was present in nine positive hits that belong to seven different pharmacological categories. To confirm that the phenylpentanol scaffold is vital for the azole chemosensitizing activity observed in these hits, we evaluated the activity of sex additional phenylpentanol derivatives that were not present in the Pharmakon library. Four additional azole chemosensitizing agents were identified, including the most structurally basic compound, 5-phenylpentanol. The finding that 5-phenylpentanol, by itself, was able to interact synergistically with fluconazole against *C. albicans* stresses the vital contribution of this scaffold to the azole chemosensitizing activities observed with phenylpentanol derivatives. Interestingly, the structurally similar scaffold phenylethanol was reported to interact synergistically with fluconazole, indicating that further optimization of these scaffolds is feasible [31].

Notably, our results indicated that the fluconazole chemosensitizing activity of pitavastatin was superior to all other tested phenylpentanol derivatives, including the pharmacologically-related statins, which have been previously reported to possess azole chemosensitizing activities [32]. Moreover, the pitavastatin-fluconazole combination displayed a broad-spectrum synergistic activity and was effective against *C. albicans, C. glabrata*, and the emerging multidrug-resistant *C. auris*. Additionally, the synergy existing between pitavastatin and fluconazole extended to interfering with the biofilm-forming ability of *C. albicans, C. glabrata*, and *C. auris*. Subinhibitory concentrations of the pitavastatin-fluconazole combination significantly compromised the ability of *Candida* species to form adherent biofilm structures more than either pitavastatin or fluconazole alone. Also, in a *C. elegans* whole animal infection model, the pitavastatin-fluconazole combination significantly reduced the burden of *C. albicans, C. glabrata*, and *C. auris* in infected nematodes. Altogether, these findings suggest the pitavastatin-fluconazole combination may have clinical implications, especially in controlling infections caused by azole-resistant *Candida* species.

Of note, the spectrum of the synergistic relationship between pitavastatin and azoles was found to be dependent on the azole drug. Pitavastatin displayed a broader spectrum of azole chemosensitizing activities when it was combined with either fluconazole or voriconazole (89% of *Candida* strains were susceptible to these combinations). However, the combination of pitavastatin and itraconazole exhibited a synergistic activity only against nine (or 50%) *Candida* strains. Although significant synergistic interactions were detected between pitavastatin and fluconazole against all the tested azole-resistant *C. auris* isolates, this was not sufficient to reduce fluconazole MICs to the susceptibility breakpoint [33]. However, the combination of pitavastatin with newer azoles completely restored the susceptibility to itraconazole and voriconazole in all the tested multidrug-resistant *C. auris*.

The azole chemosensitizing activity of statins has been attributed to their ability to interfere with the fungal ergosterol biosynthesis, however, this mechanism does not explain statins’ inconsistent effects against efflux-hyperactive *Candida* strains [28, 34]. As shown earlier, pitavastatin demonstrated a significant fluconazole chemosensitizing activity against *C. albicans* strains SC-TAC1^R34A^ and TWO7243 which have been reported as ABC-efflux hyperactive strains [35-37]. However, the pitavastatin-fluconazole combination was ineffective against *C. albicans* strains SC-MRR1^R34A^ and TWO7241 which have been reported as MFS-efflux hyperactive strains [35-37]. Moreover, the interruption of ergosterol biosynthesis often results in a fungistatic activity, which does not explain the fungicidal activity displayed by statins, as reported by others [38]. Accordingly, Lima *et al.* proposed the existence of additional mechanisms by which statins induced their antifungal activities. Interestingly, statin drugs such as simvastatin, lovastatin, and atorvastatin, have been reported to inhibit P-glycoproteins, the human analog of the fungal ABC membrane transporters [39, 40]. Therefore, we hypothesized that pitavastatin might possess the ability to interfere with *Candida*’s efflux activity through inhibition of ABC membrane transporters. To investigate this, we measured Nile Red efflux from four *C. albicans* strains known to have upregulated efflux activities. The Nile red fluorescence data revealed pitavastatin interfered specifically with efflux activities mediated by ABC membrane transporters. The finding that the fungal ABC transporters can equally respond to mammalian ABC inhibitors provides further corroboration to the existence of strong evolutionary structural conservation of these transporters [41, 42]. More interestingly, the basic 5-phenylpentanol scaffold significantly reduced Nile Red efflux in all four efflux-hyperactive *C. albicans* strains, suggesting a broad-spectrum efflux inhibitory activity by 5-phenylpentanol (Supplementary Figure-1).

In conclusion, we were able to identify phenylpentanol as a common core structure present in the most potent azole chemosensitizing agents identified in our screening. The pitavastatin-fluconazole combination exhibited broad-spectrum synergistic interactions against clinically-relevant *Candida* species. Remarkably, pitavastatin was able to restore the susceptibility of the multidrug-resistant *C. auris* isolates to the antifungal activities of itraconazole and voriconazole. The azole chemosensitization activity displayed by pitavastatin is attributed, at least in part, to its ability to interfere with *Candida’s* efflux machinery. Furthermore, the efflux inhibitory activity observed with 5-phenylpentanol provides a new scaffold for medicinal chemists to synthesize more potent and novel fungal efflux pump inhibitors.

## Materials and Methods

### Fungal strains and culture reagents

Fungal strains used in this study are listed in Supplementary Table-1. *C. albicans* clinical isolates TWO7241 and TWO7243 were obtained from professor Theodor White (UMKC). SC5314 mutant derivatives SC-MRR1^R34A^ and SC-TAC1^R34A^, containing gain-of-function alleles (MRR1P683S and TAC1G980E), were obtained from professor David Rogers (University of Tennessee Health Science Center). RPMI 1640 powder with glutamine, but without NaHCO_3_, was purchased from Thermo Fisher Scientific (Waltham, MA). 3-(N-Morpholino) propanesulfonic acid (MOPS) was obtained from Sigma Aldrich (St. Louis, MO). YPD broth medium and YPD agar were obtained from Becton, Dickinson Company (Franklin Lakes, NJ).

### Chemicals and Drugs

The Pharmakon 1600 library was purchased from MicroSource Discovery Systems, Inc. (Gaylordsville, CT). Compounds were delivered in microplates (10 mM, dissolved in DMSO) and stored at −80°C until use. Doxycycline and 5-Phenyl-1-pentanol were obtained from Alfa Aesar (Tewksbury, MA). Nile red, voriconazole, and itraconazole were obtained from TCI America (Portland, OR). Apomorphine, cholecalciferol, (R)-γ-Benzyl-L-proline hydrochloride, 5-(3, 5 Dimethylphenyl)-5-oxovaleric acid, lovastatin, and norgestimate were obtained from Sigma Aldrich (St. Louis, MO). Simvastatin, pitavastatin, and 6-Gingerol were obtained from Ark Pharm (Arlington Heights, IL). Atorvastatin was obtained from Selleckchem (Radnor, PA). Dehydroabietic acid was purchased from Pfaltz & Bauer (Waterbury, CT). Fluconazole was obtained from Fisher Scientific (Pittsburgh, PA). Gentamicin sulfate was purchased from Chem-Impex International INC. (Wood Dale, IL).

### Screening of Pharmakon library and structurally-related compounds

The Pharmakon 1600 drug library was screened against the azole-resistant *C. albicans* NR-29448. An overnight culture of *C. albicans* NR-29448 was diluted to approximately 0.5 - 2.5 × 10^3^ cells/ml in RPMI 1640 medium buffered with 0.165 M MOPS reagent. Aliquots (100 µl) of the fungal suspension was transferred to the wells of round-bottomed 96-well microtitre plates containing 16 µM concentrations of the tested drug library. The screening was conducted twice either in the presence or absence of a sub-inhibitory concentration of fluconazole (32 µg/ml). The plates were incubated at 37 °C for 24 hours. Drugs that visually reduced the growth of *C. albicans* by at least 50 % (only in the presence of fluconazole) were identified as “positive hits.”

### Microdilution checkerboard assays

The interactions between the identified hits and different antifungal drugs were assessed using broth microdilution checkerboard assays, as previously reported [43-45]. ΣFICI (fractional inhibitory concentration index) is used to measure the interaction between the tested combinations. ΣFICI interpretation corresponded to the following definitions: synergism (SYN), ΣFICI ≤ 0.5; additivity (ADD), ΣFICI > 0.5 and ≤ 1; and indifference, ΣFICI > 1 and ≤ 4 [46].

### Growth kinetics

The effect of the pitavastatin-fluconazole combination on the growth kinetics of *C. albicans* NR-29448, *C. glabrata* ATCC MYA-2950, and *C. auris* 390 were monitored using a spectrophotometer. Briefly, overnight cultures of *Candida* species were adjusted to 0.5-2.5 × 10^3^ CFU/ml in RPMI 1640 medium. Then, the pitavastatin-fluconazole combination, at the respective MIC identified from the checkerboard assays, were incubated with different *Candida* isolates at 35°C. Fungal growth was monitored by measuring the optical density at 595 nm (OD_595_) at 0, 6, 12, 18, 24, 36, and 48 h after incubation. The growth kinetics curves were constructed by plotting the OD values at the corresponding time points.

### Biofilm inhibition assay

Three *Candida* species, *C. albicans* NR-29448, *C. glabrata* HM-1123, and *C. auris* 385 demonstrated prominent ability to form robust adherent biofilms. As such, these strains were used to study the antibiofilm activity of the pitavastatin-fluconazole combination. The microtiter biofilm formation assay using crystal violet was used, as previously described [26]. Briefly, overnight cultures of the tested *Candida* strains, grown in YPD broth, were diluted in RPMI 1640 medium to approximately 1 × 10^5^ CFU/ml. Then 100 µl aliquots of each suspension were transferred to wells in 96-well plates. Aliquots of two-fold serial dilutions of pitavastatin, at concentrations ranging from 64 to 0.25 µg/ml, were transferred to the 96-well biofilm assay plates and plates were incubated at 35°C for 24 h. Pitavastatin was tested either individually or in combination with fluconazole (2 µg/ml). The adherent biofilms were then rinsed twice with phosphate-buffered saline (PBS) and left to dry at room temperature. Air-dried biofilms were stained with crystal violet (0.1%). Stained biofilms were rinsed thrice with PBS and then air dried. The resultant biofilm biomasses were quantified by dissolving the crystal violet-stained biofilms in absolute ethanol before recording absorbance values (OD_595_).

### Nile red glucose-induced efflux assay and flow cytometry

Nile Red efflux assays were performed following a previously reported protocol [30, 47, 48]. Nile red fluorescence intensity was measured at an excitation wavelength of 485 and an emission wavelength of 528 using the SpectraMax i3x microplate reader (Molecular Devices, CA, USA). Detection of fluorescence intensity was commenced about 15 seconds after glucose addition (T_0_) and then in one-minute intervals for 10 minutes. At any given time point, the fluorescence intensity was expressed as the relative change in the fluorescence intensity with regard to its initial time signal at T0. For flow cytometric analysis, starved yeast cells were incubated with Nile red (7.5 µM) for 30 minutes at 35 °C. After staining, cells were washed twice with cold PBS and then exposed to the tested treatments: glucose (40 mM), 2-deoxyglucose (40 mM), pitavastatin (0.25 × MIC), or 5-phenylpentanol (0.25 × MIC). After 30 minutes, cells were fixed in 2% paraformaldehyde and were examined in a Canto II flow cytometer (BD Bioscience, San Jose, CA), following a previously reported protocol [49]. Data were analyzed using FlowJo software v10 (Tree Star, Ashland, OR).

### *Caenorhabditis elegans* fungal infection model

To examine the *in vivo* efficacy of pitavastatin in enhancing the activity of fluconazole against azole-resistant *C. albicans*, we used the *C. elegans* animal model following previously reported guidelines [18, 26] *C. albicans* NR-29448, *C. glabrata* ATCC MYA-2950, and *C. auris* 390 displayed enhanced susceptibility to the effect of the pitavastatin-fluconazole combination and were selected for this experiment. Briefly, L4 stage worms [strain AU37 genotype glp-4(bn2) I; sek-1(km4) X] were infected by co-incubating them with approximately 5 × 10^7^ CFU/ml of *Candida* suspensions for 3 h at room temperature. After infection, worms were washed five times with M9 buffer and transferred into microcentrifuge tubes (20 worms per tube). Infected worms were treated with different combinations of pitavastatin and fluconazole at concentrations equivalent to 5, 10, or 20 µg/ml. Treatments with either PBS, pitavastatin or fluconazole alone served as controls. Worms were treated for 24 h at 25 °C. Posttreatment, worms were examined microscopically to evaluate morphological changes and ensure viability. Worms were then washed with M9 buffer five times and then disrupted by vigorous vortexing with silicon carbide particles. The resulting *Candida* suspensions were serially diluted and transferred to YPD agar plates containing gentamicin (100 g/ml). Plates were incubated for 48 h at 35°C before the viable CFU per worm was determined.

### Statistical analyses

All experiments were performed in triplicates and repeated at least three times. Statistical analyses were performed using GraphPad Prism 6.0 (Graph Pad Software, La Jolla, CA, USA). *P*-values were calculated using one-way ANOVA, and *P*-values < 0.05 were considered significant. Data are presented as means ± standard deviation.

## Acknowledgments

The fungal isolates used in this study were generously provided by BEI Resources and the US Centers for Disease Control and Prevention (CDC). The authors would like to thank Professor Theodor White (University of Missouri-Kansas City) for kindly providing *C. albicans* clinical isolates TWO7241 and TWO7243. Also, The authors would like to thank Professor David Rogers (University of Tennessee Health Science Center), for graciously providing *C. albicans* SC5314 mutant constructs (SC-TAC1^R34A^ and SC-MRR1^R34A^) used in the efflux experiments. Finally, The authors would like to thank Dr. Haroon Mohammad (Purdue University) for language editing and proofreading the manuscript.

